# Overcoming ibrutinib resistance by targeting phosphatidylinositol-3-kinase signaling in diffuse large B-cell lymphoma

**DOI:** 10.1101/523761

**Authors:** Neeraj Jain, Ondrej Havranek, Ram Kumar Singh, Tamer Khashab, Fazal Shirazi, Lalit Sehgal, Felipe Samaniego

**Affiliations:** Department of Lymphoma and Myeloma, The University of Texas MD Anderson Cancer Center, Houston, TX; Biocev, 1st Faculty of Medicine, Charles University in Prague, Czech Republic; Department of Internal Medicine, Lankenau Medical Center, Wynnewood, PA; Division of Hematology, Department of Internal Medicine, The Ohio State University, Columbus, OH

## Abstract

Diffuse large B-cell lymphoma is the most common subtype of non-Hodgkin lymphoma; 40% of patients relapse following a complete response or are refractory to therapy. The activated subtype of diffuse large B-cell lymphoma relies upon B-cell receptor signaling for survival; this signaling can be modulated by the activity of Bruton’s tyrosine kinase. Targeting that kinase with its inhibitor ibrutinib provides a potential therapeutic approach for the activated B-cell subtype of diffuse large B-cell lymphoma. However, non-Hodgkin lymphoma is often resistant to ibrutinib or soon develops resistance after exposure to it. In this study, we explored the development of acquired ibrutinib resistance. After generating three isogenic ibrutinib-resistant diffuse large B-cell lymphoma cell lines, we investigated the deregulated pathways that are associated with colony formation, growth rates, and tumorigenic properties. We found that reduced levels of Bruton’s tyrosine kinase and enhanced phosphatidylinositol 3-kinase/AKT signaling were hallmarks of these ibrutinib-resistant cells. Upregulation of phosphatidylinositol-3-kinase-beta expression in those cells drove resistance and was reversed by the blocking activity of phosphatidylinositol-3-kinase-beta/delta. Treatment with the selective phosphatidylinositol-3-kinase-beta/delta dual inhibitor KA2237 reduced both tumorigenic properties and survival-based phosphatidylinositol-3-kinase/AKT/mTOR signaling of these ibrutinib-resistant cells. Additionally, combining KA2237 with currently available chemotherapeutic agents synergistically inhibited the metabolic growth of these ibrutinib-resistant cells. This study elucidates the compensatory upregulated phosphatidylinositol-3-kinase/AKT axis that emerges in ibrutinib-resistant cells.

## Introduction

Diffuse large B-cell lymphoma (DLBCL), the most common subtype of non-Hodgkin lymphoma (NHL), accounts for ~30% of total NHLs.^1^ While curable with R-CHOP treatment in the majority of DLBCL patients, up-to one third of those patients develop relapsed/refractory disease, a major cause of mortality and morbidity.^2, 3^ DLBCL, a heterogeneous lymphoma, can be classified into major molecular subtypes (activated B-cell [ABC] and germinal center B-cell (GCB]) based on distinct gene expression and genetically mutational signatures.^4^ Importantly, compared to GCB-DLBCL patients, the ABC population has lower survival rates following multi-agent chemotherapy.^5, 6^ Since ABC-DLBCL is characterized by chronically active B-cell receptor (BCR) signaling, several components of BCR signaling pathways are emerging as attractive therapeutic targets.^4^ Bruton’s tyrosine kinase (BTK), the critical component of BCR signaling, drives the BCR signaling cascade leading to activation of NF-κB and other target signaling.^7, 8^ Ibrutinib, an orally administered BTK inhibitor, has been FDA-approved to treat patients with relapsed mantle cell lymphoma, Waldenstrom’s macroglobulinemia, and chronic lymphocytic leukemia, including those harboring the 17p deletion.^9, 10^ In a phase I/II clinical trial of relapsed/refractory DLBCL, ibrutinib treatment resulted in an overall response rate of 37% in ABC-DLBCL patients versus 5% in GCB-DLBCL patients, indicating that the ABC subtype is more susceptible to BTK targeting.^4^ Despite these encouraging results, responses to ibrutinib treatment are variable/incomplete and show drug-resistance and population/genetic alterations stemming from unknown causes.^11, 12^

BCR signaling, initiated by self-antigen reactivity of BCR or by mutation in MYD88, activates both NF-κB in the ABC-DLBCL survival pathway and the phosphatidylinositol-3-kinase (PI3K) signaling pathway.^7, 13, 14^ The class-I subPI3K family includes the alpha-, beta-, gamma-, and delta isoforms, which are often constitutively activated in cancer.^15^ Koo et al reported that pan-PI3K inhibitors, which target all PI3K isoforms, exert a reduction in cell viability in a subset of ABC-DLBCL lines with CD79 mutations.^13^ However, due to broad toxicities of pan-PI3K inhibitors, attention has shifted to use of single PI3K isoform specific inhibitors to treat cancer.^16^ Idelalisib, a PI3K-delta specific inhibitor, received FDA approval for treatment of B-cell malignancies.^17–19^ On the other hand, inhibition of PI3K-delta in ABC-DLBCL cells led to activation of PI3K-alpha via a compensatory mechanism, defeating treatment intent.^20, 21^

We have identified PI3K-beta/delta-mediated activation of AKT as a compensatory survival pathway that is potentially responsible for the emergence of ibrutinib-related resistance in ABC-DLBCL cells. Treatment of ibrutinib resistant (IR) DLBCL cell lines with a selective dual PI3K-beta/delta inhibitor significantly reduced the AKT activity and tumor volume in xenografts. Moreover, when combined with currently used chemotherapeutic agents, the PI3K-beta/delta inhibitor strongly inhibited the growth of IR-DLBCL cells. This combination could provide an additional therapeutic strategy for overcoming ibrutinib resistance in DLCBLs.

## Methods

### Cell culture and drugs

ABC-DLBCL cell lines (TMD8, U2932, and HBL1) and GCB lines (SU-DHL-6 and SU-DHL-8) were maintained in RPMI-1640 media supplemented with 10% fetal bovine serum. OCI ABC and -GCB lines (OCI-LY1, OCI-LY3, OCI-LY7, OCI-LY8, and OCI-LY10) were maintained in Iscove's Modified Dulbecco's Medium with 20% human serum. All cell lines were regularly tested for mycoplasma and identity confirmation by Short Tandem Repeat. The BTK inhibitor ibrutinib (PCI-32765) and PI3K isoform specific inhibitors PI3K-alpha (alpelisib), PI3K-beta (AZD6482), PI3K-delta (idelalisib), and PI3K-alpha/delta (pictilisib) were purchased from Selleckchem. The PI3K-beta/delta dual inhibitor (KA2237) was provided by Karus Therapeutics Ltd. (Oxfordshire, UK).

### Generation of IR ABC-DLBCL cell lines

IR-DLBCL cell lines (HBL1-Ib^R^, TMD8-Ib^R^, and OCI-LY10-Ib^R^) were generated by continuously culturing parental lines (HBL1, TMD8, and OCI-LY10) with incremental doses of ibrutinib. The IC_50_s of the IR cell lines were 1200 nmol/L, 500 nmol/L, and 1740 nmol/L for OCI-LY10-Ib^R^, TMD8-Ib^R^, and HBL1-Ib^R^, respectively.

Stable knockdown of the PI3K-beta isoform in IR-DLBCL cells was performed using lentiviruses expressing human shRNA (Dharmacon; clone IDs V3LHS_341250 and 341251) as described previously.^22^

### Cell viability, colony formation, and apoptosis assay

To quantify surviving and/or proliferating cells, an MTT cell proliferation assay kit (Promega Corporation, Madison, WI, USA) was used according to the manufacturer’s instructions. In each experiment, the average relative absorption (OD_490_-OD_700_) was used to estimate the number of metabolically active cells. The percent of treated cells that survived and normalized, compared to control cells, was calculated to evaluate cell viability. A colony formation assay was performed using a methylcellulose medium (H4100; STEMCELL Technologies Inc.) as described previously.^23^ Synergistic combination indexes (CIs) between two drugs were calculated using MTT cell proliferation assays performed with combinations of different drug concentrations and using Calcusyn software. A CI value less than or equal to 0.8 indicates synergistic effects between two drug combinations.

For apoptosis assays, cells were washed with cold phosphate-buffered saline, fixed in 70% ethanol, and stained with a 0.5 µM propidium iodide solution (BD Pharmingen, BD Biosciences, San Jose, CA, USA). Propidium iodide staining was detected with flow cytometry (LSR Fortessa flow cytometer; BD Biosciences, San Jose, CA, USA), and the subG1 fraction was presented as the apoptotic population.

### Xenograft study

Animal studies were completed under the Institutional Animal Care and Use Committee (IACUC)-approved protocol for animal welfare at The MD Anderson Cancer Center, Houston, TX. Twenty-eight nude mice were subcutaneously injected with 5 × 10^6^ OCI-LY10 cells (parental-or IR-DLBCL) in a suspension containing 1:1 Matrigel (Corning, Life Sciences). When the tumors reached approximately 300 mm^3^, mice in each group were randomly assigned (7 mice to each group) to treatment every other day by intraperitoneal injection with either KA2237 (100 mg/kg) or saline control. Tumor volumes (1/2 [length X width^2^]) were measured thrice weekly.

### Statistical analysis

Results were presented as means ± standard deviation of triplicates. Statistical analyses were carried out using the Student’s *t*-test with P values less than 0.05 considered to be significant.

## Results

### Acquired ibrutinib resistance also enhances tumorigenic properties of ABC-DLBCL cells

To understand the mechanism of ibrutinib resistance in DLBCL cells, we generated isogenic, resistant versions of 3 ABC-DLBCL cell lines (HBL1, TMD8, and OCI-LY10). The three lines were selected out of ten tested based on their sensitivity to ibrutinib (*Online Supplementary Figure S1 A-C)*; note that all GCB-DLBCL cell lines examined in this study were resistant to ibrutinib. For these tests, the parental cell lines underwent *in vitro* culturing with incremental doses of ibrutinib (Figure 1A and *Online Supplementary Figure S1D*). Compared to parental cells, IR-DLBCL cells (HBL1-Ib^R^, TMD8-Ib^R^, and OCI-LY10-Ib^R^) had higher proliferation rates (Figure 1B) and exhibited lower apoptosis signaling upon ibrutinib challenge (*Online Supplementary Figure S1 E-F)*. Acquiring ibrutinib resistance was associated with increased colony number when plated in methylcellulose substrate (Figure 1C).

**Figure 1.**
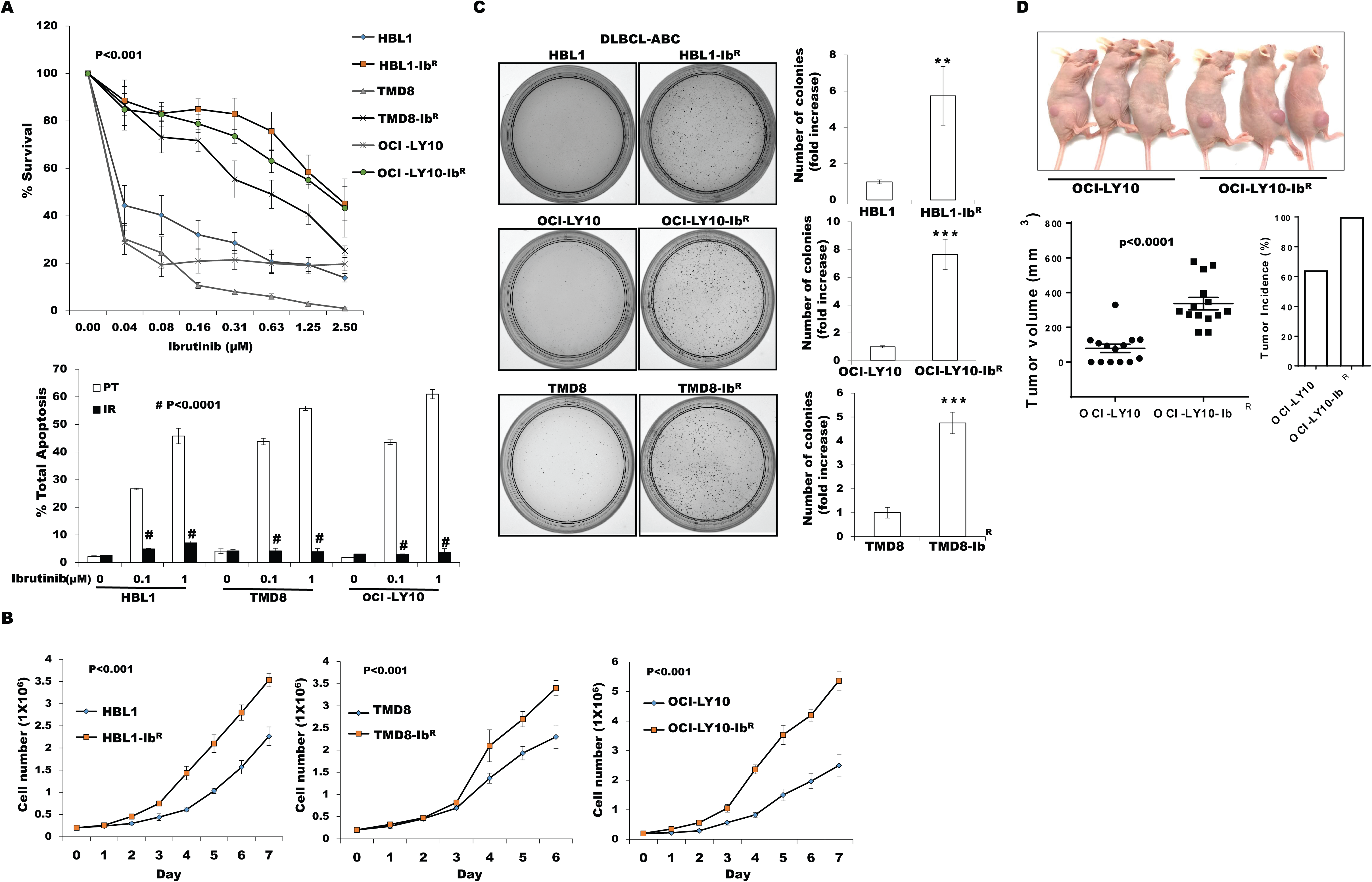
Generation of acquired ibrutinib-resistant DLBCL cells and clonogenic properties. (A) Ibrutinib-resistant (IR) DLBCL cell lines were generated from parental (PT) lines as described in Methods. The percentage of cell survival and total apoptosis were determined after 72 hours of ibrutinib treatment. (B) DLBCL cell growth rates were determined by proliferation assay; we counted cells for 7 days and compared to growth rates of respective PT lines. (C) Representative images and quantification data from colony formation assays (** *p*<0.01, *** *p*<0.001). (D) OCI-LY10 tumor cells (PT/IR) suspended in a 1:1 Matrigel mixture were implanted subcutaneously into nude mice. Twenty days after injection, tumor volumes were measured. Compared to the PT-group, tumors were larger in the IR-DLBCL group (*P*<0.0001).

Furthermore, we tested the tumorigenic potential of IR-DLBCL cells and found that those cells in nude mice xenografts had a higher tumor incidence and larger tumors compared to parental DLBCLs (parental, 9/14; IR, 14/14; P=0.04, Fisher’s exact test) (Figure 1D), indicating that IR-DLBCL cells are clearly tumorigenic.

### Cells with acquired resistance ibrutinib (IR-DLBCL) show upregulated PI3K/AKT/mTOR signaling

We identified reduced BTK expression in cultured IR-DLBCL cell lines (Figure 2A) similar to the reduced BTK expression acquired by chronic lymphocytic leukemia patients following ibrutinib treatment.^24^ BTK mutation C481S and others (e.g., PLCγ2) have been reported to be associated with ibrutinib resistance.^25, 26^ However, in the IR-DLBCL cell lines examined in this study, targeted sequencing for BTK and PLCγ2 genes confirmed the lack of these mutations (data not shown). We anticipated that dysregulated signaling pathways could lead to development of ibrutinib resistance so we assayed the protein expression in our parental- and IR-DLBCL pairs by reverse phase protein array (RPPA). The RPPA data shown by heat maps (Figure 2B) revealed that 118 proteins were differentially expressed (±1.5-fold changes in expression). We observed the upregulation of DNA damage repair signaling molecules (CHK1, CHK2, and CDCD25C) in IR-DLBCL cells compared to parental cells; these results were further validated by Western blots (data not shown). However, treatment with CHK1 or CHK2 inhibitors alone or in combination with ibrutinib did not sensitize the IR-DLBCL cells (data not shown). From our RPPA data, we also identified enhanced PI3K/AKT/mTOR signaling in IR-DLBCL—an increase in expression of its activator IRS1 and a decrease in expression of its negative regulator PTEN. We further validated this activation of PI3K/AKT signaling by Western blot analyses of paired IR/parental DLBCL cells which confirmed enhanced levels of pAKT, p-mTOR, and the latter’s substrate p-P70S6K in all three IR-DLBCL cell lines examined (Figure 2C).

**Figure 2.**
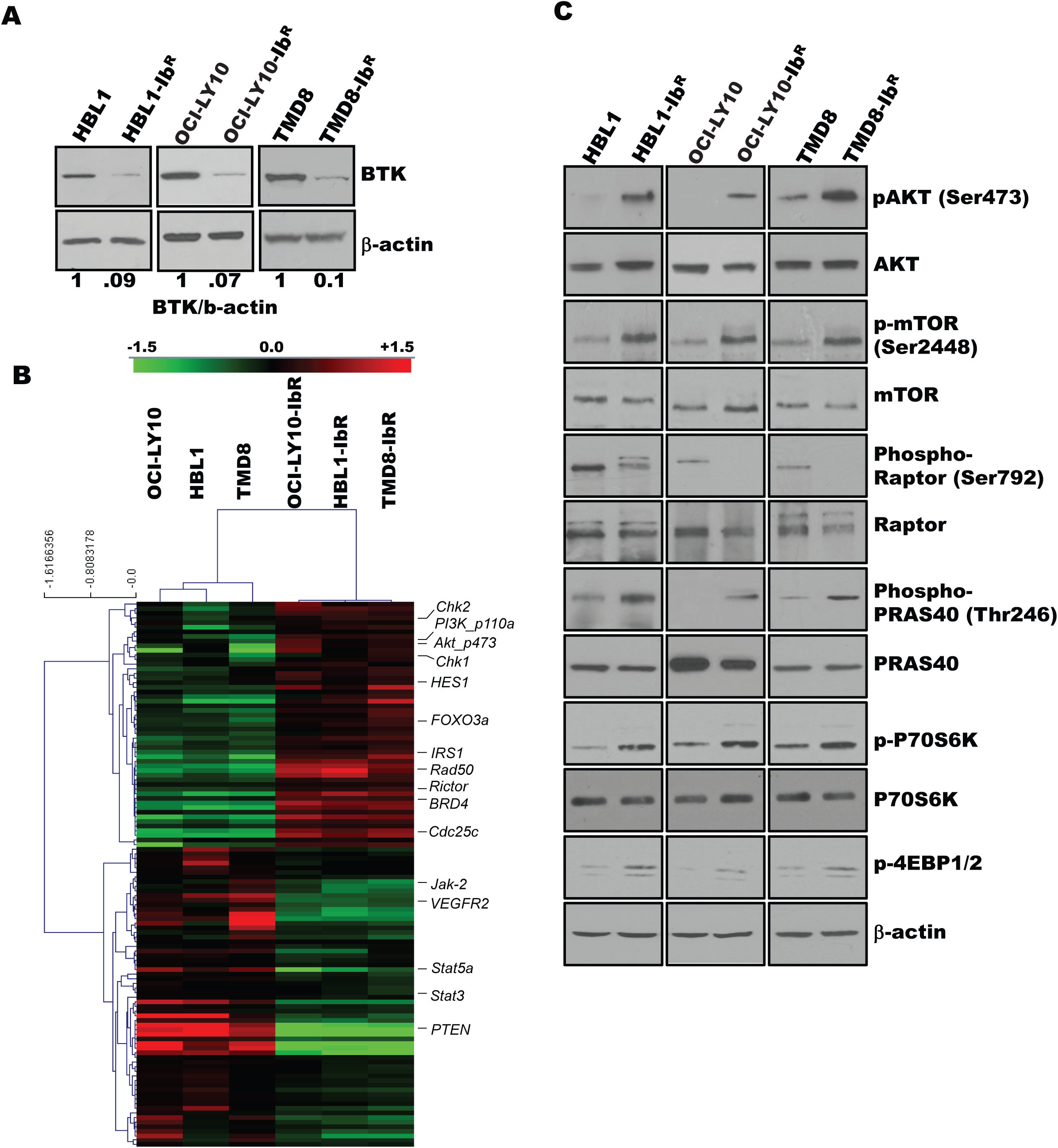
Ibrutinib-resistant (IR) DLBCL cells show enhanced PI3K/AKT/mTOR signaling. (A) Western blot analyses of BTK expression in IR/parental (PT) DLBCL cells. (B) Heat maps derived from reverse-phase protein array analyses of IR/PT DLBCLs pairs represent the differential expression (up- and downregulation) of proteins identified by Wilcoxon Rank sum test (map: red color indicates above median, green-indicates below median). (C) Western blot analyses for AKT, and mTOR and its substrates show activity in IR/PT DLBCL cell pairs.

### Simultaneous inhibition of PI3K-beta and delta isoforms sensitized IR-DLBCL cells

We identified upregulated PI3K/AKT/mTOR signaling in IR-DLBCL cells (Figure 2), and then investigated how this signaling affects ibrutinib resistance. The PI3K pathway is constitutively deregulated in multiple malignancies; studies support the importance of PI3K signaling in leukemias and lymphomas.^15, 27^ Loss of PTEN is a well-described mechanism of upregulation of PI3K/AKT/mTOR but loss of PTEN does not offer a practical restorative therapeutic option and thus we did not pursue this topic.^28^ A previous study indicated the importance of PI3K signaling in the development of ibrutinib resistance in a patient-derived xenograft model but a possible mechanism was not proposed.^29^ A recent report of mantle cell lymphoma demonstrated that inhibition of the PI3K-alpha isoform in the tumor microenvironment overcame ibrutinib resistance.^30^ Because pan-PI3K inhibitors might induce off-target cellular toxicity, we elected to examine the effects of various pharmacological inhibitors of PI3K with different isoform inhibition profiles. We first screened the expression levels of PI3K isoforms in our IR/parental DLBCL pairs; we observed a significant increase in the PI3K-alpha and -beta levels (~2- and 2.5-fold, respectively), a decrease in the PI3K-delta levels (~half-fold change), and no change in the PI3K-gamma levels (Figure 3A). We then hypothesized that enhanced PI3K-alpha or -beta isoform levels might be responsible for overall upregulated PI3K/AKT/mTOR signaling and the survival of our IR-DLBCL cells. Therefore, we treated those cells with PI3K isoform-specific inhibitors and found that PI3K-alpha (alpelisib), -beta (AZD6482), and -delta (idelalisib) responded marginally as single agents in an MTT cell proliferation assay and that they triggered no significant changes to pAKT levels (Figure 3 B-C).

**Figure 3.**
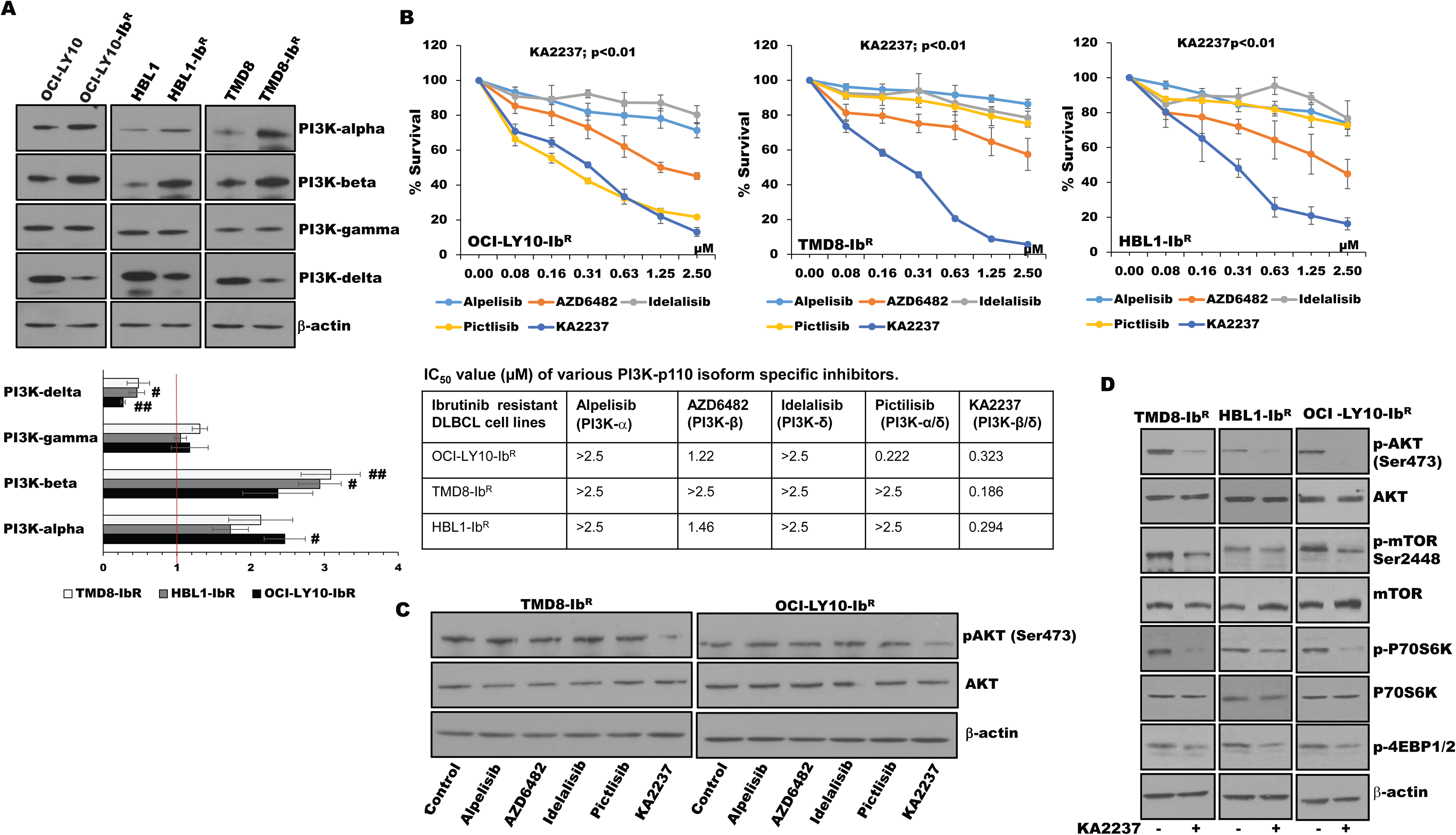
PI3K inhibitors kill ibrutinib-resistant (IR) DLBCL cells. (A) Western blots and quantification of class-I subPI3K isoforms in IR/parental (PT) DLBCL cells. (B) Metabolic activity analysis for IR-DLBCL cells treated with specific inhibitors PI3K-alpha (alpilisib), PI3K-beta (AZD6482), PI3K-delta (idelalisib), PI3K-alpha/delta (pictlisib), or PI3K-beta/delta (KA2237). The table shows IC_50_ values (µM) of inhibitors tested in IR-DLBCL cells. (C) pAKT levels in IR-DLBCL cells after 24 hours of treatment with the indicated inhibitors. (D) Western blots representing pAKT levels and activity of mTOR substrate in IR-DLBCL cells after treatment with the PI3K-beta/delta inhibitor.

We speculated that the lack of effect with these single inhibitors might be due to secondary compensatory mechanisms or feedback effects through activation of other PI3K isoforms, since a previous report noted that inhibition of PI3K-delta caused feedback activation of PI3K-alpha in the ABC subtype of DLBCL.^21^ After treatment with a dual PI3K-alpha/delta inhibitor (pictlisib), our IR-DLBCL model did not show significant changes in pAKT levels or sensitization (Figure 3 B-C). However, treatment with the selective dual PI3K-beta/delta inhibitor KA2237 significantly reduced IR-DLBCL viability and blocked the activation of the AKT/mTOR axis (Figure 3D) suggesting that PI3K-beta and -delta are the key nodes with AKT/mTOR axis signaling and cell survival in IR-DLBCL.

### Targeting the PI3K-beta isoform sensitized IR-DLBCL cells to a PI3K-delta inhibitor

To understand the potential role of the PI3K-beta isoform in the development of ibrutinib resistance, we performed a stable knockdown of that isoform in our IR-DLBCL cells using two independent shRNAs; neither caused changes in expression of other PI3K isoforms (Figure 4A). We noted that the PI3K-beta-isoform knockdown did not alter the colonogenic ability or cell growth rate of IR-DLBCL cells (data not shown). PI3K-beta knockdown did not change the activity of the AKT/mTOR axis (Figure 4C) nor did it sensitize cells to ibrutinib (data not shown). However, consistent with our results using a PI3K-beta/delta dual inhibitor (Figure 3), treatment of the PI3K-beta-knockdown cells with the PI3K-delta inhibitor idelalisib uniformly reduced the growth rates of IR-DLBCL cells (Figure 4B) and significantly reduced AKT/mTOR activity in all three IR-DLBCL cell lines (Figure 4 B-C) suggesting a PI3K-delta-mediated activation of compensatory survival signaling in PI3K-beta-knockdown IR-DLBCL cells. We further treated IR-DLBCL cells with PI3K-beta and -delta specific inhibitors administered separately and in combination. Treatment with either agent alone did not reduce cellular viability, but when combined, the treatments acted synergistically and significantly reduced cellular viability (Figure 4D and *Online Supplementary Figure S2* and *Table S1)*. In both cell types, the loss of cell viability was preceded by reduced pAKT levels (data not shown).

**Figure 4.**
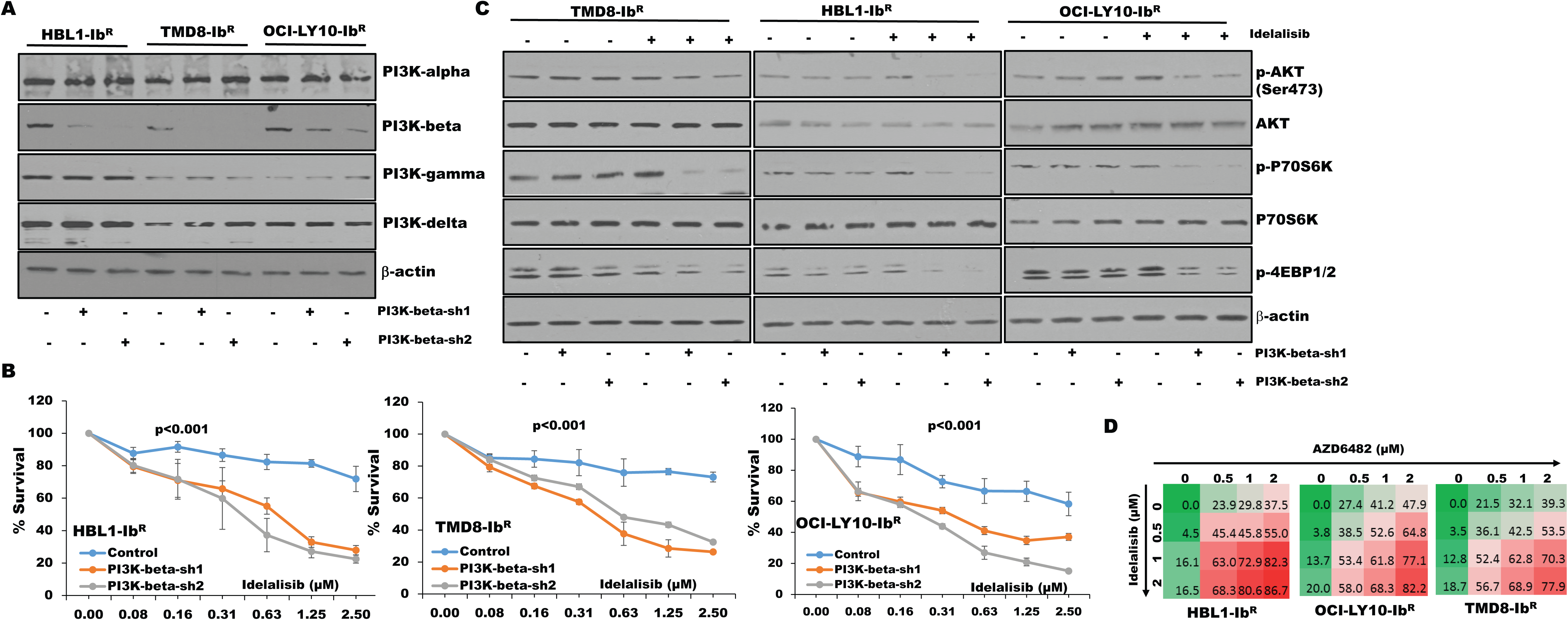
Enhanced blocking with combined PI3K-beta and -delta targeting in ibrutinib-resistant (IR) DLBCL cells. (A) Knockdown of PI3K-beta expression was performed in IR-DLBCL cell lines by stable expression of PI3K-beta isoform-specific shRNA as described in Methods. Expression analyses of PI3K isoforms demonstrate the success of PI3K-beta knockdown in IR-DLBCL cell lines. (B) Viability of cells treated with idelalisib after PI3K-beta isoform knockdown of IR-DLBCL cells was analyzed using an MTT cell proliferation assay. (C) Western blots representing pAKT levels and activity of mTOR substrate in PI3K-beta knockdown IR-DLBCL cells after idelalisib treatment. (D) Drug dose matrix data of IR-DLBCL cells. The numbers in the matrix indicate the percentage of growth inhibition of cells treated with inhibitors (AZD6482 and idelalisib) either as single agents or in combination, relative to vehicle-control treated cells. Data were visualized over the matrix using a color scale.

### Combining a PI3K-beta/delta dual inhibitor with chemotherapeutic agents sensitized IR-DLBCL cells

Given the potent activity of the selective dual PI3K-beta/delta inhibitor KA2237, we explored its effects on tumorigenic properties. As a single agent, KA2237 exerted a robust reduction in the colony-formation ability of our IR-DLBCL cells (Figure 5A and *Online Supplementary Figure S3A*); and in a xenograft model, the agent (100mg/kg, intraperitoneal, every other day) inhibited the growth of OCI-LY10-Ib^R^ DLBCL tumors (Figure 5B). Analysis of tumor extracts showed that KA2237 significantly reduced pAKT levels and concomitantly decreased mTOR activity (Figure 5C). We then explored the potential therapeutic application of this PI3K-beta/delta inhibitor when used with standard DLBCL chemotherapeutic agents. We treated our IR-DLBCL cells with KA2237 in combination with each of three such agents (doxorubicin, vincristine, or vorinostat) and observed synergy (*Online Supplementary Table S2)* and a significant reduction in cell growth (Figure 5D and *Online Supplementary Figure S3B*).

**Figure 5.**
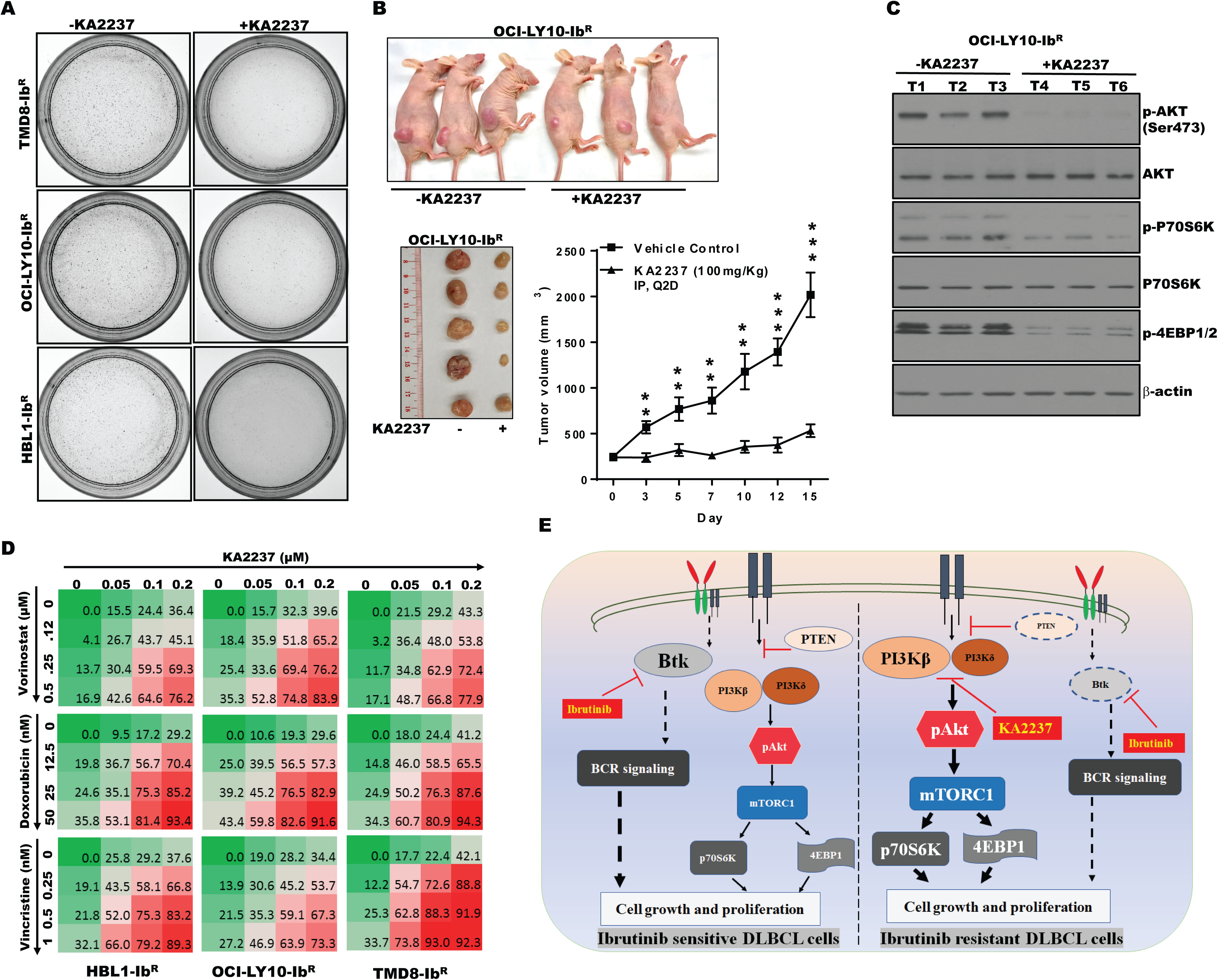
PI3K-beta/delta dual inhibitor sensitized ibrutinib-resistant (IR) cells to cytotoxic effects of chemotherapeutic agents. (A) Colony formation assays performed using IR-DLBCL cells treated with KA2237. (B) OCI-LY10-Ib^R^ tumor cells suspended in a 1:1 Matrigel mixture were implanted subcutaneously into nude mice. Intraperitoneal administration of either KA2237 (100 mg/kg) or saline control was initiated every other day after tumors reached ~300 mm^3^. Tumor volumes were reported for all mice for 15 days (** *p*<0.01, *** *p*<0.001). (C) Western blot analysis for PI3K/AKT/mTOR signaling pathway from tumor lysates treated with KA2237. (D) Drug-dose matrix data of respective IR cells. The numbers in each matrix indicate the percent of growth inhibition, relative to vehicle-alone cells, of cells treated with KA2237 plus one of three chemotherapeutic agents. (E) Graphical image representing the PI3K-beta/delta-dependent activation of survival PI3K/AKT signaling in acquired IR DLBCL cells. This activation of survival PI3K signaling is blunted by the dual PI3K-beta/delta selective inhibitor KA2237, which has a heightened effect when combined with currently used chemotherapeutic agents. BCR, B-cell receptor; BTK, Bruton’s tyrosine kinase. Note: font sizes of the molecules in this figure represent their experimentally observed expression levels.

## Discussion

Aggressive lymphomas such as ABC-DLBCL are a major cause of morbidity and mortality, primarily due to the development of therapeutic resistance.^6^ Chronic activation of BCR signaling by BTK provides survival benefits to ABC-DLBCL cells; thus, BCR signaling remains an important therapeutic target for patients who are refractory to, or have relapsed from, current R-CHOP chemotherapy.^31, 32^ The FDA-approved BTK inhibitor ibrutinib has shown significant efficacy in treating ABC-DLBCL patients; however ibrutinib resistance frequently follows.^7, 11, 32^ Using lines examined in this study, we discovered that ibrutinib-resistance is acquired through compensatory activation of PI3K/AKT/mTOR signaling, which supports survival and enhances the tumorigenic properties in these IR ABC-DLBCLs.

Furthermore, we found that PI3K/AKT signaling is maintained in our IR-DLBCL cell lines through the PI3K-beta and delta isoforms. We confirmed the dependency of these IR ABC-DLBCL cells on the PI3K-beta/delta subunits by knocking down the PI3K-beta isoform in acquired IR-DLBCL cell lines and by inhibiting the PI3K-beta/delta isoforms that effectively blocked PI3K/AKT signaling and induced death of IR-DLBCL cells.

Previous reports suggest that patients who are nonresponsive to ibrutinib may often have mutations in BTK (BTKC481S) and/or PLCγ2 (R665W,L845F,S707Y).^33^ However, our targeted sequencing of three IR-DLBCL cell lines identified no such genetic alterations in these two genes (data not shown) suggesting that the selection of mutant clones during generation of IR lines is not a cause/mechanism of acquired resistance. Another study confirmed the role of mutant MYD88 expression in providing partial resistance to ibrutinib in ABC-DLBCL cell lines.^34^ We observed the overexpression of BCL2 and inhibitor-of-apoptosis family members (cIAP, xIAP, and survivin) in our IR-DLBCLs cells (*Online Supplementary Figure S1F*). This is consistent with a previous report that identified elevated BCL2 expression following development of ibrutinib resistance.^35^ While venetoclax, an FDA-approved BCL2 inhibitor, can promote anti-proliferative activity for IR-DLBCL, targeting BCL2 could lead to the development of resistance to a BCL2-inhibitor. As an example, one study showed that either short- or long-term exposure of B-cell lymphoma cell lines (MCL and DLBCL) to venetoclax led to the development of venetoclax resistance due to a reduction in PTEN expression (a negative regulator of the PI3K/AKT activation pathway).^36^ Our RPPA analysis from three IR cell lines showed a significant decrease in PTEN expression with enhanced expression of activated AKT (p-AKT) and downstream activation of mTOR signaling, suggesting the critical role of PI3K signaling in survival of acquired IR-DLBCLs (Figure 2C).

Given the high frequency of PI3K-pathway activation in human cancers, several PI3K inhibitors are being tested in clinical trials. This includes pan-PI3K inhibitors, which demonstrate great efficacy but also produce adverse toxicity effects.^16^ As such, selective PI3K isoform specific inhibitors are of interest. The PI3K-delta specific inhibitor idelalisib was approved by the FDA for treating relapsed chronic lymphocytic leukemias and B-cell lymphomas, yet the drug has not provided significant improvement in overall survival.^19, 37-39^ Multiple studies have revealed the feedback activation of PI3K pathways or compensatory effects among various PI3K isoforms in response to PI3K inhibitors. For example, activation of PI3K-beta isoform was identified in preclinical studies after PI3K-alpha was blocked.^40–42^ Therefore, targeting multiple PI3K isoforms with dual PI3K-isoform specific inhibitors might eliminate compensatory/feedback responses caused by targeting single PI3K-isoform selective drugs. In support, one study showed tumor regression using a PI3K-alpha/delta dual inhibitor (copanlisib) in CD79B(WT)/MYD88(mut) patient-derived IR ABC-DLBCL cells, underscoring a genetic mechanism of ibrutinib resistance regulated by PI3K isoforms.^20^ Our acquired IR-DLBCL model did not show significant changes in pAKT levels or sensitization after treatment with a dual PI3K-alpha/delta inhibitor (pictlisib) even though we observed a significant increase in the PI3K-alpha isoform (Figure 3 B-C). We speculated that this was due to differential activation of the signaling pathways between the derived genetic mutations and the chronically-exposed, acquired-IR model. Along with an increase in PI3K-alpha expression, we identified a strong and significant increase in expression of the PI3K-beta isoform in our acquired IR-DLBCL model (Figure 3A). Treatment with the single PI3K-beta specific inhibitor (AZD6482) did not alter the pAKT level nor kill the IR-DLBCL cells (Figure 3 B-C); whereas, treatment with a PI3K-beta/delta dual inhibitor (KA2237) significantly reduced AKT/mTOR signaling and profoundly reduced IR xenograft tumors (Figures 3 and 5). To note, we also observed reduced expression of the PI3K-delta isoform in IR-DLBCL cells compared to parental cells, suggesting that even modest expression of the delta isoform might be mediating feedback activation of PI3K signaling in IR-DLBCL cells. After knocking down a PI3K-beta isoform and treating it with a PI3K-delta isoform inhibitor, we confirmed that PI3K-beta and -delta isoforms were involved in maintaining ibrutinib resistance in ABC-DLBCL cells (Figure 4). These results suggest that enhanced PI3K-beta expression might be masking the effect of the PI3K-delta inhibitor in IR lines. We observed a reduction in the PI3K-beta isoform (without expression changes in other PI3K isoforms) following treatment with the PI3K-beta/delta inhibitor KA2237 in IR cells (data not shown). This, further supports our hypothesis that enhanced expression of the PI3K-beta isoform might reduce the anti-proliferative activity of the PI3K-delta specific inhibitor in IR lines. We further substantiated our ibrutinib findings using a second targeting agent, ACP-196, a more selective inhibitor of BTK.^43^ We generated an ACP-196-resistant model of two ABC-DLBCL cell lines (TMD8 and OCI-LY10) and observed upregulation of PI3K/AKT/mTOR signaling, with sensitization of ACP-196 resistant DLBCL cells after challenge with the PI3K-beta/delta dual inhibitor (data not shown).

In summary, our results show that after gaining ibrutinib resistance, ABC-DLBCL cells have reduced expression of BTK and are dependent on PI3K/AKT/mTOR signaling for their survival. We found that upregulation of the PI3K/AKT axis was due to an increased expression of the PI3K-beta isoform, which masked the sensitization of cells to a PI3K-delta inhibitor. Targeting the PI3K/AKT/mTOR axis with a PI3K-beta/delta selective inhibitor reduced both tumorigenic properties and survival-based PI3K/AKT/mTOR signaling of ibrutinib-resistant cells. These effects were enhanced when combined with chemotherapeutic agents active against DLBCL (Figure 5E).

## Acknowledgments

The authors would like to thank the staff in the Department of Lymphoma and Myeloma at MD Anderson, including Stephanie Martch for her assistance with writing and editing the manuscript. NJ, LS, and F Samaniego, and LS designed the research studies, analyzed the data, and wrote the manuscript. NJ, RKS, TK, OH, and F Shirazi performed experiments and analyzed the data. All authors contributed critical review of the manuscript and approved the final version.

